# CED-5/CED-12 (DOCK/ELMO) can promote and inhibit F-actin formation via distinct motifs that target different GTPases

**DOI:** 10.1101/2023.10.04.560868

**Authors:** Thejasvi Venkatachalam, Sushma Mannimala, Martha C. Soto

## Abstract

Coordinated activation and inhibition of F-actin supports the movements of morphogenesis. Understanding the proteins that regulate F-actin is important, since these proteins are mis-regulated in diseases like cancer. Our studies of *C. elegans* embryonic epidermal morphogenesis identified the GTPase CED-10/Rac1 as an essential activator of F-actin. However, we need to identify the GEF, or Guanine-nucleotide Exchange Factor, that activates CED-10/Rac1 during embryonic cell migrations. The two-component GEF, CED-5/CED-12, is known to activate CED-10/Rac1 to promote cell movements that result in the engulfment of dying cells during embryogenesis, and a later cell migration of the larval Distal Tip Cell. It is believed that CED-5/CED-12 powers cellular movements of corpse engulfment and DTC migration by promoting F-actin formation. Therefore, we tested if CED-5/CED-12 was involved in embryonic migrations, and got a contradictory result. CED-5/CED-12 definitely support embryonic migrations, since their loss led to embryos that died due to failed epidermal cell migrations. However, CED-5/CED-12 inhibited F-actin in the migrating epidermis, the opposite of what was expected for a CED-10 GEF. To address how CED-12/CED-5 could have two opposing effects on F-actin, during corpse engulfment and cell migration, we investigated if CED-12 harbors GAP (GTPase Activating Protein) functions. A candidate GAP region in CED-12 faces away from the CED-5 GEF catalytic region. Mutating a candidate catalytic Arginine in the CED-12 GAP region (R537A) altered the epidermal cell migration function, and not the corpse engulfment function. A candidate GEF region on CED-5 faces towards Rac1/CED-10. Mutating Serine-Arginine in CED-5/DOCK predicted to bind and stabilize Rac1 for catalysis, resulted in loss of both ventral enclosure and corpse engulfment. Genetic and expression studies showed the GEF and GAP functions act on different GTPases. Thus, we propose CED-5/CED-12 support the cycling of multiple GTPases, by using distinct domains, to both promote and inhibit F-actin nucleation.

**Author Summary:** GTPases in their active state promote actin nucleation that drives cellular events, from cell migrations, to cell shape changes, to cell-cell interactions. To function correctly, GTPases need to cycle from the active, GTP-bound state, to the inactive, GDP-bound state. This cycle is supported by Guanine-nucleotide Exchange Factors, or GEFs, that support activation as GDP is switched for GTP, and GTPase-Activating Proteins, or GAPs that support hydrolysis back to the GDP bound state. The Rac1/CED-10 GTPase has a well-studied GEF, CED-5/CED-12, that promotes Rac1 activation during cell engulfment of dying cells. Here we tested if CED-5/CED-12 also functioned as the activator, or GEF for Rac1 during embryonic cell migrations. Surprisingly, CED-5/CED-12 behaved completely opposite to what was expected during this cell migration. Therefore, we investigated if CED-5/CED-12 could harbor a GAP function. Comparing models of human and *C. elegans* protein structures suggested a putative GAP region, which we mutated to show that CED-12 likely functions as a GAP. Genetic and gene expression tests identify other GTPases, CDC-42 and RHO-1, regulated by this newly uncovered CED-12 GAP function. This places CED-5/CED-12 at a central position, where it can support the cycling of multiple GTPases, and both promote and inhibit F-actin nucleation.

## INTRODUCTION

Organized F-actin is required for cell migrations, and for engulfment of dying cells and cellular debris. The idea that the processes of cell migration and cell/corpse or debris engulfment use related actin regulators has been proposed for processes like macrophage migration and engulfment in Drosophila (Wood and Davidson, 2020 Cell Reports). The idea that embryonic cell migrations and cell corpse engulfment are linked in *C. elegans* was suggested when we cloned mutations in the Arp2/3 regulators, *gex-2* and *gex-3*, and reported loss of these genes resulted in both long-lived corpses and epidermal migration defects, similar to loss of *Rac1/ced-10* (Soto et al., 2002). These studies raised the question of how actin is regulated during two apparently distinct processes, epidermal enclosure, a collective cell migration by the epidermal cells to engulf the entire embryo, and corpse engulfment, where part of the membrane of individual cells reorganizes to engulf their dying neighbors.

Epiboly, the process that creates the metazoan body plan, depends on regulated sheet migrations. In *C. elegans*, epiboly begins at the 400-cell stage, when the main embryonic tissues begin to differentiate, and the epidermis begins its complex migrations. An open question in all organisms, is how signals are coordinated in tissues to promote the correctly oriented movements. By focusing on the first movements of the epidermal cell sheet, ventral enclosure and dorsal intercalation, we have identified a pathway that promotes polarized enrichment of branched actin: the GTPase CED-10/Rac1 activates the WAVE/Scar complex, a nucleation promoting factor (NPF) for Arp2/3. Our studies identified signals that localize CED-10 and WVE-1 correctly at membranes, to direct F-actin enrichment (Bernadskaya et al., 2012). However, how these membrane signals are communicated to the CED-10 GTPase during epiboly is not understood. The guanine exchange factors, or GEFs, which help to activate GTPases by promoting the transition from GDP to GTP (Rossman et al., 2002; Bos et al, 2007), are excellent candidates to connect signals at the membrane to the activation of CED-10/Rac1. However, the CED-10 GEF or GEFs that promote embryonic epidermal movements are not known.

Mutations in CED-10/Rac1 were first identified for their role in cell corpse engulfment (Ellis et al., 1991). The original *ced-10* alleles were hypomorphic alleles that interfered with the ability of cells to engulf dying cells (Ellis et al., 1991; Reddien and Horvitz 2000). Identifying null alleles of *ced-10* revealed a second role for CED-10: promoting embryonic morphogenesis (Lundquist et al., 2001; Soto et al., 2002). CED-10 was proposed to recruit and activate the WAVE complex, which in *C. elegans* is essential for embryonic organ tissue formation and tissue migrations (Soto et al., 2002; Patel et al., 2008). GTPases are proposed to be recruited and activated at membranes by guanine-nucleotide exchange factors, GEFs, that enhance the exchange of GDP for GTP, thus activating the GTPase (Reviewed in Rossman et al., 2005; Bos et al 2007). The GEF for CED-10 during corpse engulfment was proposed to be CED-5/DOCK180 (Reddien and Horvitz 2001). CED-5 was also proposed to act as the GEF for CED-10 during neuronal migrations (Stavoe and Colon-Ramos 2012).

CED-5/DOCK180 is a DOCK GEF which requires an ELMO protein to activate Rac (Brugnera et al., 2002). In *C. elegans*, CED-5/DOCK180 works with CED-12/ELMO to activate CED-10 during corpse clearance (Proposed by Conradt 2001; Gumienny et al. 2001; Wu et al, 2001; Chung et al, 2000; Zhou et al., 2001). During corpse engulfment CED-5/CED-12 and CED-10 have been proposed to act in neighboring cells, to enclose dying cells, by promoting F-actin polymerization in membrane protrusions that encircle dying cells. F-actin was shown to enrich around dying cells in the germline and embryos (Kinchen et al., 2005; Shen et al, 2013), and then disappear as the corpse decayed. While some studies noted that CED-12 appears to have an embryonic role (Gumienny et al, 2001), the role of CED-5 and CED-12 during embryonic development, and epidermal morphogenesis has not been investigated.

We present here analysis of the role of CED-5/DOCK and CED-12/ELMO, during epidermal morphogenesis, that shows these cell death regulators also support epidermal migrations. We therefore investigated if CED-5/CED-12, a candidate GEF for CED-10’s ventral enclosure function, shares epidermal *ced-10* phenotypes. While loss of the candidate GEF, CED-5/CED-12 reduced F-actin around corpses, as expected, loss of CED-5/CED-12 increased F-actin in epidermal cells during ventral enclosure, the opposite phenotype as loss of CED-10. To determine how a GEF can both promote and inhibit F-actin formation, we investigated if this candidate CED-10/Rac1 GEF also functions as a RhoGAP. A candidate GAP domain was identified in CED-12 and mutated. Loss of the GAP function of CED-12 resembled loss of CED-12 for epidermal migrations, but corpse engulfment was normal. GEF function specific mutations were created by CRISPR, to test if these affected both processes. These studies identify a previously undescribed role during embryonic morphogenesis for CED-5 and CED-12, proteins well-studied for their cell corpse engulfment. Analysis of a mutation in the newly identified GAP domain of CED-12 suggest CED-12/ELMO, uses different subdomains to switch from promoting to inhibiting F-actin formation, depending on the subcellular context. Other GTPase targets of the CED-5/CED-12 epidermal morphogenesis function are identified, placing DOCK/ELMO at a central position for actin regulation during morphogenesis.

## RESULTS

### CED-5/DOCK and CED-12/ELMO regulate morphogenesis by modulating F-actin in migrating epidermal cells

CED-5/DOCK-180 and CED-12/ELMO act as a bipartite GEF that promotes active CED-10/Rac1 during the engulfment of cells that die by programmed cell death in *C. elegans* (Chung et al, 2000; Gumienny et al 2001; Zhou et al., 2001; Wu et al, 2001). We showed that CED-10/Rac1 also acts to promote the tissue migration known as ventral enclosure that leads the epidermis to enclose the developing *C. elegans* embryo (Soto et al. 2002; Fig. 1A). However, the GEF for CED-10/Rac1 during this process is not known. To test if CED-5 and CED-12 are required during embryonic epidermal cell migrations, we crossed null or strong loss of function alleles of *ced-5(n1812)* (Wu and Horvitz 1998) and *ced-12(n3261) (*Zhou et al., 2001) into a strain that expresses F-actin only in the epidermis (*lin-26p::Lifeact::mCherry*, Havrylenko et al., 2015). Staged embryos that were oriented with ventral tissues facing up (towards the cover slip) were imaged from 280 to 320 minutes after first cleavage. These 4D movies of live embryos showed that wild type embryos are enriched for F-actin at the leading edge of the migrating row of ventral cells, as previously shown (Bernadskaya et al, 2012; Raduwan et al., 2020). Timing the migration, from the first appearance of leading cell edge protrusions, to when the leading cells met at the ventral midline, took approximately 20 minutes at 23 C. Loss of the GTPase CED-10/Rac1, showed significantly reduced levels of F-actin in leading cells, and arrested migrations, as would be expected for the removal of a major activator of branched actin (Fig. 1B-D). If CED-5/CED-12 are the GEF that activates CED-10 during ventral enclosure, we expected removing them would similarly lead to decreased F-actin levels and slower or arrested migrations. Surprisingly, animals carrying null mutations in *ced-5* or *ced-12* displayed two opposite phenotypes to loss of CED-10. First, the levels of F-actin in the leading cells were significantly higher than in controls (Fig. 1B). Second, the ventral cells met at the midline significantly faster, on average 5 minutes, or 25% faster (Fig. 1C). We obtained the same results with multiple null or strong loss of function alleles, including *ced-5(n1812, n2002), ced-12(n3261, ky149)*, and when *ced-5* was removed by RNAi. Therefore, the candidate GEF for CED-10 during ventral enclosure had the opposite effect on F-actin levels and migration timing as was expected.

**Fig. 1:**
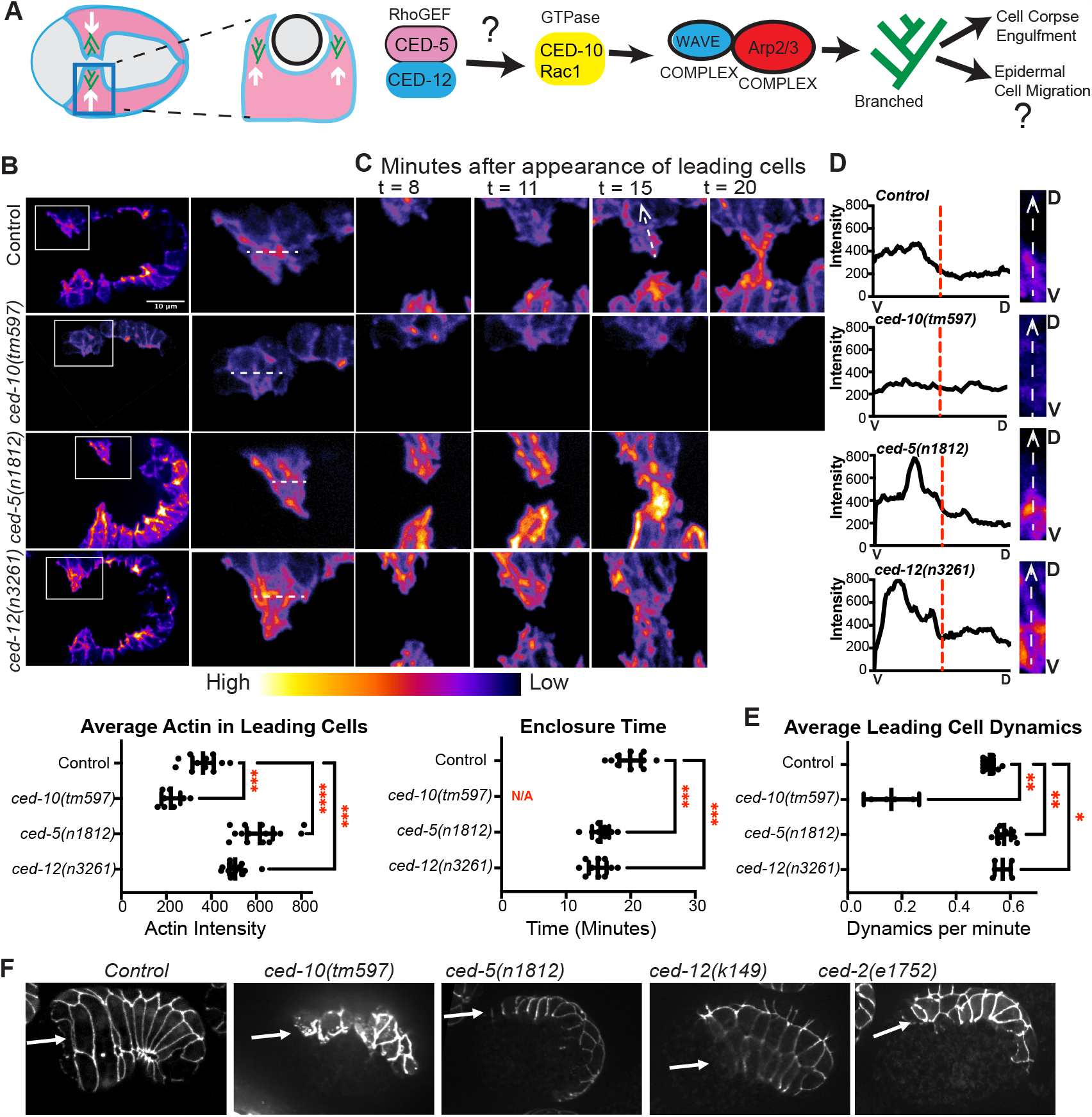
CED-5/CED-12 DOCK/ELMO regulate morphogenesis. A. Cartoon introducing ventral enclosure, a collective epidermal cell migration essential for embryonic development, and cell corpse engulfment, a movement in the same epidermal cells. The proteins proposed to regulate branched actin to regulate these movements are shown: the GEF CED-5/CED-12 (DOCK/ELMO) activates CED-10/Rac1, which activates the WAVE complex, a nucleation promoting factor for Arp2/3 complex. (B) Epidermal F-actin in live embryos was imaged using the *lin-26p::LifeAct::mCherry* strain (REF). Embryos were oriented ventral up and anterior to the left. Leading cells (LCs) were boxed and magnified. White line across LC shows how actin intensity was measured, and added to the plot below. Each dot represents the average of three measurements. n= at least 12 embryos. (C) Comparison of the time it takes for the LCs to meet, showing selected time points after their first appearance (t = 8, 11, 15, and 20 minutes). Enclosure time, plotted below, was measured from the first appearance of LCs. *ced-10(tm597)* was marked as N/A as LCs never meet. (D) Representative line scans illustrate distribution of actin intensity from the ventral half compared with the dorsal half of LCs. Close-ups of representative Leading Cells, V= ventral (bottom), D=dorsal (top). A ventral to dorsal line was drawn, as shown in the t15 Control, dotted white arrow, and intensity was plotted using Fiji’s Plot Profile tool. Dashed red lines indicate half the cell’s length. (E) Dynamics (protrusions plus retractions) of LCs during enclosure were were summed up from LCs’ first appearance to time of leading cells meeting, divided by total time, and plotted as dynamics per minute. (F) Morphogenesis defects visualized by the *dlg-1::gfp (xnIs16)* transgene that marks the junctions of epithelial tissues. White arrows point to same row of ventral epidermal cells in the wild type and mutant embryos. Ventral cells in mutant embryos fail to migrate, leading to Gex (<u>g</u>ut on the <u>ex</u>terior) phenotype (Soto et al., 2002). Asterisks mark statistical significance, *=p<0.05, **=p<0.01, ***=p<0.001, ****=p<0.0001, as determined by one-way Welch’s ANOVA test. In this and all Figures error bars show 95% confidence intervals.

Proper migration of the embryonic epidermis requires dynamic F-actin at the leading edge, that is and ventrally enriched (Bernadskaya et al., 2012). The *ced-5* and *ced-12* mutants showed properly polarized, ventrally enriched F-actin (Fig. 1D). To test if the higher actin levels, and faster migrations correlated with increased dynamics at the leading edge of migrating cells, we measured protrusions and retractions. While a null mutation in *ced-10, tm597*, led to strongly decreased protrusions and retractions, null mutations in *ced-5* and *ced-12* resulted in significantly increased protrusions and retractions (Fig. 1E). Elevated F-actin and more rapidly moving cells may disrupt the migrations. If this was the case, we expected to see altered behavior in the migrating epidermal cells.

### Changes in F-actin levels and cell migration speed contribute to morphogenesis defects

Since loss of CED-10, and of its proposed GEF CED-5/CED-12 caused opposite effects on epidermal F-actin levels, we next examined the consequences of the mutations for embryonic morphogenesis. When proteins are removed that contribute to the epidermal cell migrations, it leads to the Gex, or <u>g</u>ut on the <u>ex</u>terior phenotype (Soto et al. 2002), which can be seen with differential interference contrast (DIC) optics, and by following the epithelial apical junctions using transgenic strains like FT48 *xnIs16 dlg-1::gfp* (Totong et al., 2007). We compared the frequency of embryonic lethality with a Gex phenotype, for embryos missing *ced-10, ced-5, ced-12* or the adaptor *ced-2/CRKII*, which is thought to act with *ced-5* and *ced-12* to support corpse engulfment *(Hedgecock et al., 1983; Ellis et al., 1991; Reddien and Horvitz, 2000; Chung et al., 2000; Kinchen et al., 2005)*. Complete loss of *ced-10* is as severe as complete loss of the WAVE complex, with 100% of the embryos dying with a Gex phenotype. Partial reduction mutations in *ced-10*, like *n1993*, a mutation in the CED-10 prenylation site, thought to reduce membrane attachment (Reddien and Horvitz 2000), results in partial Gex lethality, of approximately 15%. Loss of *ced-5* or *ced-12* using putative null alleles led to partial Gex lethality (14%, 14-16% lethality), more similar to the partial loss of function *ced-10* alleles (Fig. 1F, Table 1). Mutations in *ced-2* had a similar and somewhat milder phenotype (9% lethality, Fig. 1F and Table 1). These results reinforced that high levels of F-actin can interfere with morphogenesis, as we previously showed for loss of *vab-1* (Bernadkaya et al, 2012), *hum-7* (Wallace et al., 2018) and *rga-8* (Raduwan et al., 2020).

**Table 1.**
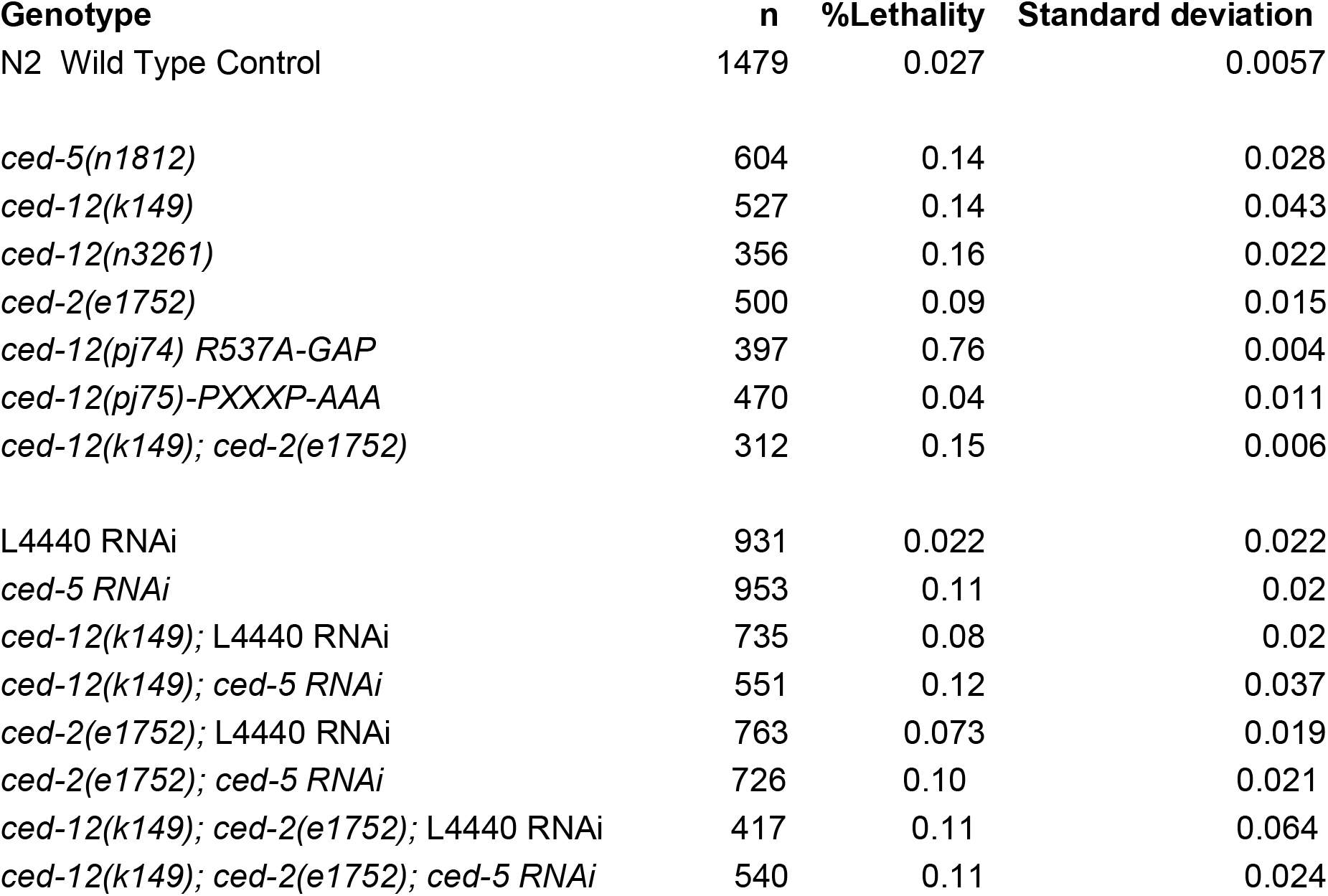
Embryonic lethality count of *ced-5, ced-12* and *ced-2* alleles.

### Embryo shape changes support that CED-5 and CED-12 function in early embryos

Since the embryonic role of CED-5 and CED-12 has not been investigated, we looked for further evidence that CED-5 and CED-12 are required in embryos. We noted that some embryos that die due to arrested epidermal cell migrations also displayed a round shape instead of the wild type oval shape (Fig. 2A). Embryo shape is established just after fertilization: unfertilized oocytes are round, and become oval after fertilization, as sperm entry is coupled to release of chitin and other egg shell components (McCarter et al., 1999; L’Hernault Wormbook 2006; Johnston et al., 2010; Johnston and Dennis, 2011; Gonzalez et al., 2018). Mutations that alter constriction of the spermatheca during fertilization result in round or otherwise misshapen embryos (Kovacevic and Cram, 2010; Tan and Zaidel-Bar, 2015). Mutations in WAVE complex components like *gex-3*, or null alleles of *ced-10* are frequently round (our unpublished observations). To compare the “round” vs. oval wild-type phenotypes, we measured the lengths and widths of embryos, and documented if the embryo lived or died. In Wild Type, the average length was 49.3um and the average width was 31.7um. Comparing wild type with *ced-5 (n1812), ced-12(n3261)* and *gex-3(RNAi)* showed that lengths were significantly shorter than wild type in the majority of mutant embryos, with the exception of *ced-12(n3261)* viable embryos (Fig. 2A,B). By comparison, though the widths were slightly increased in some mutants, the differences were not significant (Fig. 2C), suggesting the main differences in mutant embryos are in their lengths. In wild-type embryos, the ratio of length divided by width averaged 1.6, while mutations in *ced-5 (n1812), ced-12(n3261)* and *gex-3(RNAi)* resulted in significantly reduced ratios for all *ced-5(n1812)* and *gex-3 RNAi* embryos, and for *ced-12(n3261)* embryos that died (Fig. 2D).

**Fig. 2:**
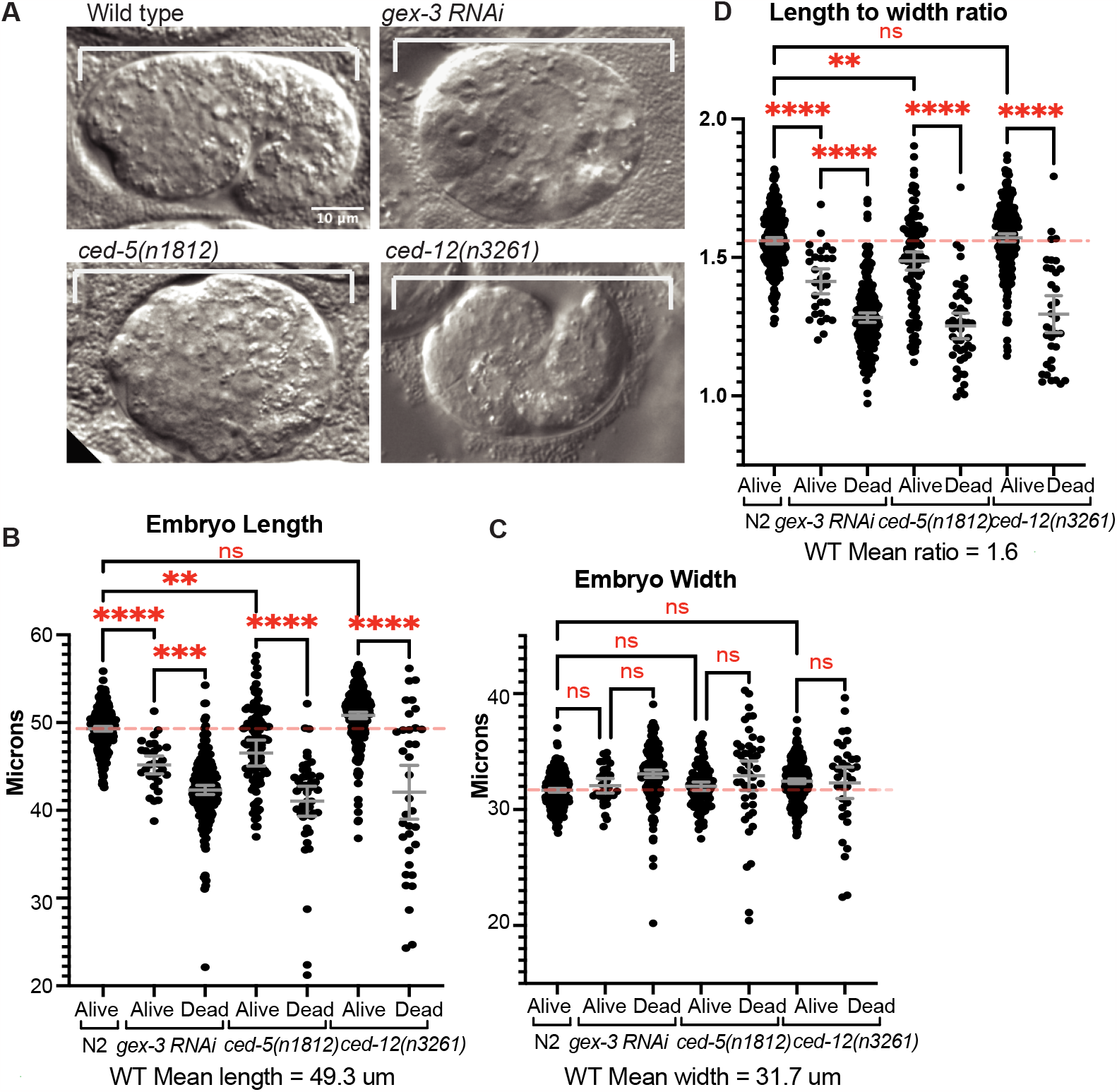
Mutant embryos that die are shorter and rounder than those that live. A. Representative differential interference contrast (DIC) images of embryos, oriented anterior to the left, and dorsal up, to illustrate a live Control embryo (Wild Type) and dying, short round embryos with reduced *gex-3, ced-5* or *ced-12*. B. Embryo length was measured in Live and Dead embryos for each genotype. C. Embryo width was measured in Live and Dead embryos for each genotype. D. The ratio of length divided by width was plotted for each genotype. Asterisks mark statistical significance, *=p<0.05, **=p<0.01, ***=p<0.001, ****=p<0.0001, as determined by one-way Welch’s ANOVA test.

Since the loss of *ced-5 (n1812), ced-12(n3261)* and *gex-3(RNAi)* resulted in a mixture of live and dead embryos, we compared how length correlated with viability. For all three mutations, dead embryos were significantly more likely to be “round” or short. In Wild Type, <1% of embryos are “round” or short, and <1% of embryos die. *ced-5 (n1812)* and *gex-3(RNAi)* embryos that lived were significantly shorter compared to wild type. However, *ced-5 (n1812), ced-12(n3261)* and *gex-3(RNAi)* embryos that died were much shorter than wild type, and significantly shorter than their sibling embryos that lived. The ratio of length divided by width dropped from 1.6 in wild type to ∼1.3 in dying mutants (Fig. 2D). Altogether these embryo shape changes suggested loss of *gex-3, ced-5* or *ced-12* affected early events of embryonic development, when embryo shape is first established.

### CED-5/DOCK, CED-12/ELMO and CED-2/CRKII together support embryonic viability

The CRKII homolog, CED-2, is part of the corpse engulfment pathway that includes CED-5 and CED-12. To test if *ced-5, ced-12* or *ced-2* work in a complex during ventral enclosure as they do during corpse engulfment, we constructed double mutants between them. All double mutants had similar levels of embryonic lethality as the single mutants, with the Gex phenotype. To test the triple mutant, we depleted *ced-5* via RNAi in animals with the *ced-12; ced-2* double mutation (Table 1). These animals had similar levels of lethality, with Gex-like morphogenesis phenotypes supporting that the three proteins also act together during ventral enclosure (Table 1).

We used a genetic test to measure if *ced-5* behaved like a *ced-10* GEF. We predicted that loss of a GEF would strongly enhance partial loss of a GTPase. Combining two hypomorphic alleles of the GTPase *ced-10 (n1993 and n3246)* with depletion of *ced-5* via RNAi resulted in no significant change in embryonic lethality (Table 2). Thus, CED-5 did not behave like a candidate GEF for CED-10/Rac1 ventral enclosure function.

**Table 2.** Effect of *ced-5* null alleles on partial loss of function *ced-10* alleles.

| Genotype | n | %Lethality | Standard deviation |
| --- | --- | --- | --- |
| *ced-10(n1993)* – *L4440* RNAi control | 318 | 11 |  |
| *ced-5* RNAi | 541 | 9 | 0.4 |
| *ced-10(n1993); ced-5* RNAi | 268 | 12 |  |
| *ced-10(n3246); ced-5* RNAi | 286 | 13 | 0.7 |

### CED-5/CED-12 and GEX-3 promote F-actin at corpses

To test if CED-10, and its proposed GEF CED-5/CED-12, have similar or opposite effects on F-actin around corpses, we crossed putative null mutations in *ced-5* or *ced-12* into a strain with labeled epidermal F-actin (*lin-26p::LA::mCherry*, used in Fig. 1) and which also expressed the receptor, CED-1, around corpses (*Pced-1::ced-1::gfp*, Zhou et al., 2001). These movies were used to measure F-actin levels around well-studied corpses, C1 (left), C2 (right) and C3 (left) in the ventral epidermis (Shen et al., 2013), that are engulfed by the same migrating epidermal cells (Fig. 3A,B). Loss of *gex-3*, an effector of CED-10 with cell migration and corpse engulfment defects (Soto et al., 2002), resulted in reduced F-actin levels around the corpses, and did loss of *ced-5(n1812)*, or *ced-12(n3261)*. Since all of the mutants have long-lived corpses that last longer than in wild type controls, this result agrees with the model that F-actin enrichment around corpses is promoted by a molecular pathway that includes the GTPase CED-10, its effector the WAVE complex, and its GEF CED-5/CED-12, to engulf and remove corpses during development (Reddien and Horivtz, 2004, Kinchen et al., 2005; Fig. 3C). Thus, monitoring F-actin in embryonic epidermal cells showed two opposing results: during corpse engulfment, loss of CED-5/CED-12 caused the expected loss of F-actin enrichment around corpses (Fig. 3), while during ventral engulfment (Fig. 1), loss of CED-5/CED-12 caused the opposite effect, increased F-actin, in the same tissue (Fig. 3C).

**Fig. 3.**
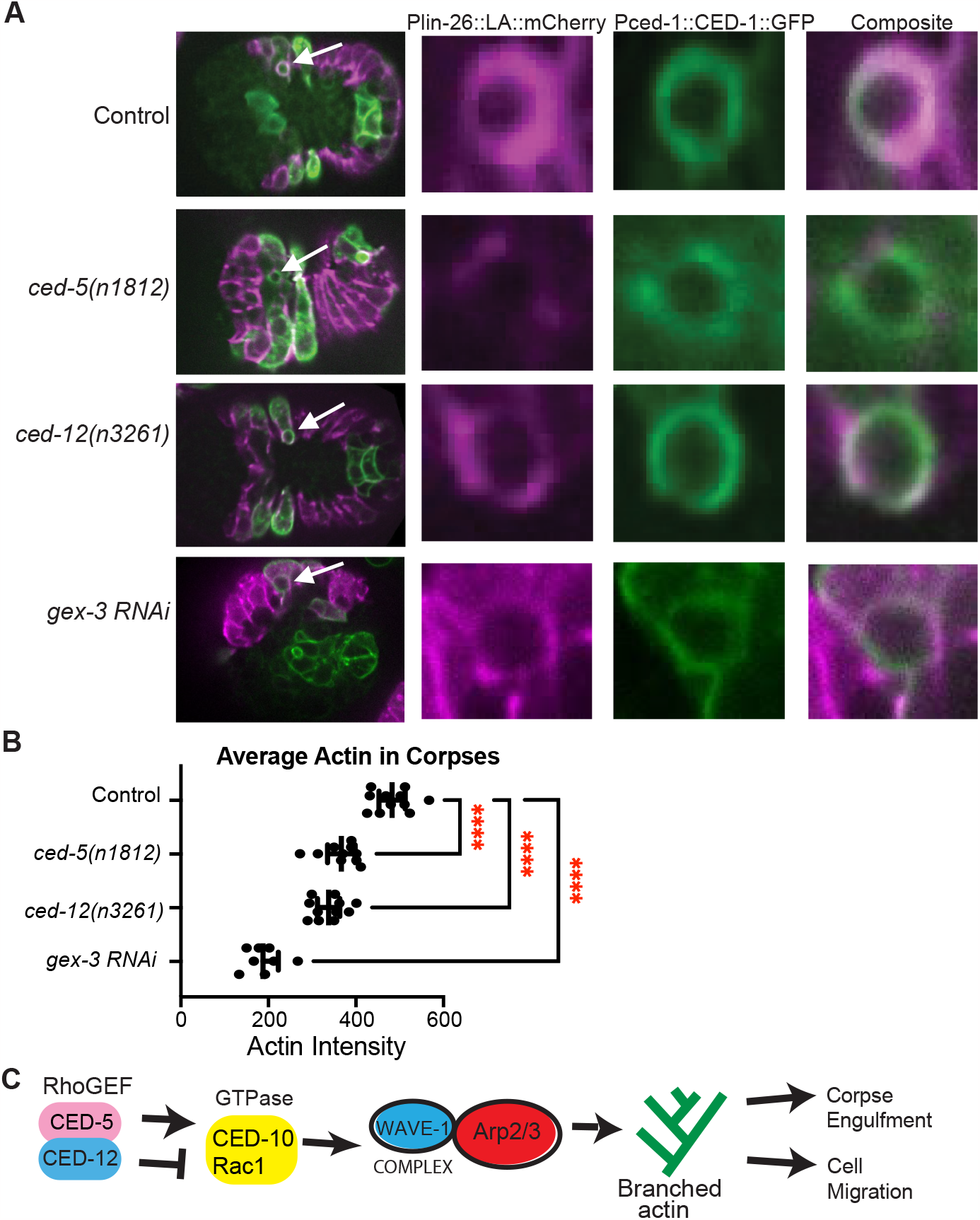
CED-5 and CED-12 promote F-actin around corpses. (A) To visualize corpse engulfment by epidermal cells, embryos with epidermal F-actin (*lin-26p::LifeAct::mCherry)*, and with the corpse receptor (*ced-1p::ced-1::gf)* were imaged live, at the same times in development. Left panels: composite image of the full embryo, white arrows indicate C1 corpse. The right panels zoom in on C1 corpse, F-actin in magenta, CED-1::GFP in green. (B) The average intensity of F-actin around C1, C2, and C3 corpses was measured, n = at least 20 cells. Asterisks mark statistical significance, ****=p<0.0001, as determined by one-way Welch’s ANOVA test. (C) CED-5/CED-12 are proposed to have two different roles regulating F-actin during two branched-actin dependent events.

### CED-12 Alignments and structural comparison identify candidate GAP motif

To address how one complex, the CED-5/CED-12 GEF, could promote two distinct and opposite functions, we looked for evidence of alternative forms of CED-5 or CED-12. The current molecular model for CED-5 predicts a single protein, and the model for CED-12 predicts two proteins that differ only by 7 amino acids at the N terminus (*wormbase*.*org –* Davis et al., 2022). While CED-12 is the only ELMO protein in *C. elegans*, other organisms, including humans and Dictyostelium, have six ELMOs, including some thought to promote F-actin formation, and others thought to inhibit F-actin. Inhibitory ELMOs, like ELMO-A in Dictyostelium and ELMO-D1 in humans may use a GAP domain (Ivanova et al., 2014). Alignments of CED-12 with *C. elegans* GAP proteins revealed a region with high homology to GAPs (Fig. 4A). Further, aligning CED-12 with ELMO and ELMOD proteins identified a similar region between CED-12 and ELMOD proteins, but not ELMO proteins (Fig. 4A). Both alignments suggested R537 of CED-12 may be the catalytic Arginine in a CED-12 GAP domain. We aligned, or threaded, the recently published structure of DOCK2/ELMO1 with active Rac (Chang et al., 2020), with *C. elegans* CED-5/CED-12 with active CED-10 using the program SWISS-MODEL and visualized them using UCSF Chimera (https://swissmodel.expasy.org/; Waterhouse et al., 2018; Goddard et al, 2018; Fig. 4B). We also aligned the closed structure (Chang et al., 2020) of DOCK2/ELMO1 with CED-5/CED-12 (Fig. 4D). The aligned structures showed that the catalytic Arginine of the proposed CED-12 GAP, R537 faces out, away from DOCK/CED-5 and Rac1/CED-10, and away from the membrane, in the DOCK2/ELMO1 structure in both the open and closed conformations (Fig. 4B-D). Thus *C. elegans* CED-12 can be fit onto the structure of the human ELMO, yet it contains a GAP motif, not found in human ELMOs, that points away from the pocket of CED-5 that binds to CED-10. Human ELMO has Q541 in place of R527 (Fig. 4B).

**Fig. 4:**
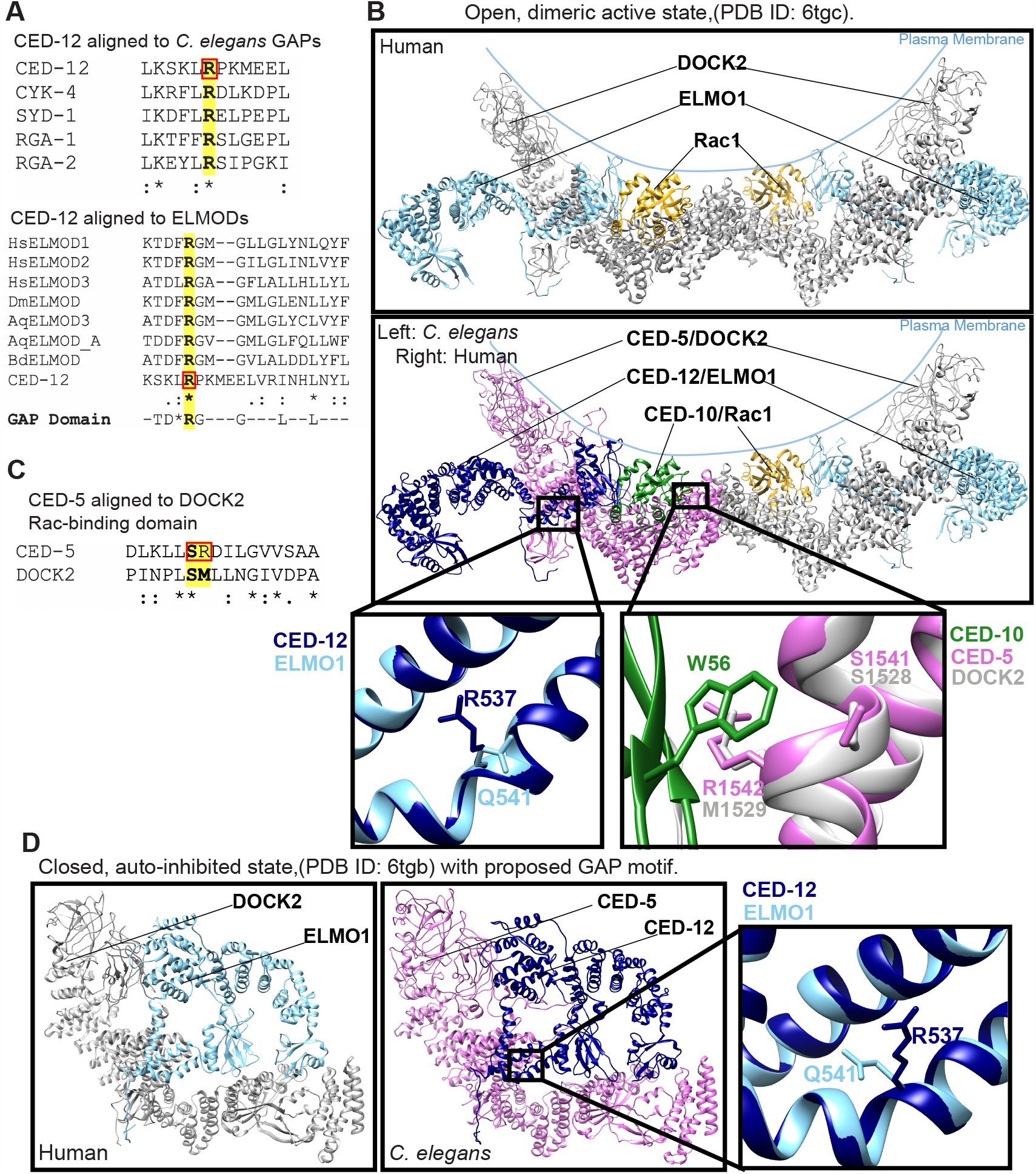
Protein alignments and models identify residues for proposed GAP and GEF functions of CED-5/CED-12. (A) Aligning CED-12 to the *C. elegans* GAPs CYK-4, SYD-1, RGA-1 and RGA-2 (top) identifies a GAP region in CED-12. The catalytic arginine of the GAPs and CED-12 is shown in boldface and highlighted. Aligning CED-12 against ELMODs from Hs (human), Dm (Drosophila), Aq (Amphimedon queenslandica) or Bd (Batrachochytrium dendrobatidis) (bottom) illustrated they have a conserved region, that includes the arginine residue R537, boxed in red in both alignments. (B) Ribbon model of open, dimeric active conformation of DOCK2/ELMO1/Rac1 and CED-5/CED-12/CED-10 (PDB ID: 6TGC) (Chang et al., 2020). The top panel depicts the human structure. The panel below shows the threaded structures for *C. elegans* CED-5/CED-12/CED-10 on the left dimerized with the human homologs on the right. Gray/pink: CED-5/ DOCK2, Light blue/navy blue: CED-12/ ELMO1, yellow/green: CED-10/ Rac1. R537 is novel to *C. elegans*, with a Q541 in the human homolog. In the Rac binding domain (RBD), CED-5 S1541/R1542 align closely to DOCK2 S1528/M1529, enabling close contacts with CED-10 W56/ Rac1 W56. (C) CED-5 aligned to the Rac-binding domain (RBD) of human DOCK2. Key residues, highlighted and in boldface, S1528/ M1529 (human) align to S1541/ R1542 of *C. elegans*. (D) Closed, auto-inhibited conformation of DOCK2/ELMO1 (left) and threaded models of *C. elegans* CED-5 and CED-12 on right (PDB ID: 6TGB). Boxed and magnified region shows novel CED-12 R537 superimposed on human ELMO1 Q541.

### CED-12 GAP (R537A) and CED-5 GEF/RBD (SR1541/1542AA) mutations affect embryonic development

If the predicted GAP motif detected in CED-12 is important for function, mutating it may indicate how this proposed GAP function supports CED-12 activities (Fig. 5A). GAP function requires a catalytic Arginine (Barrett et al., 1997; Rittinger et al. 1997). We used CRISPR to mutate the R537 to Alanine, predicted to eliminate catalytic GAP function (Wallace et al., 2018). We first examined the *ced-12(pj74)* R537A allele for embryonic lethality. We found that 7.6% of embryos died, with Gex morphogenesis phenotypes, compared to 14-16% for two putative null alleles of *ced-12* (Table 1, Fig. 5B).

**Fig. 5.**
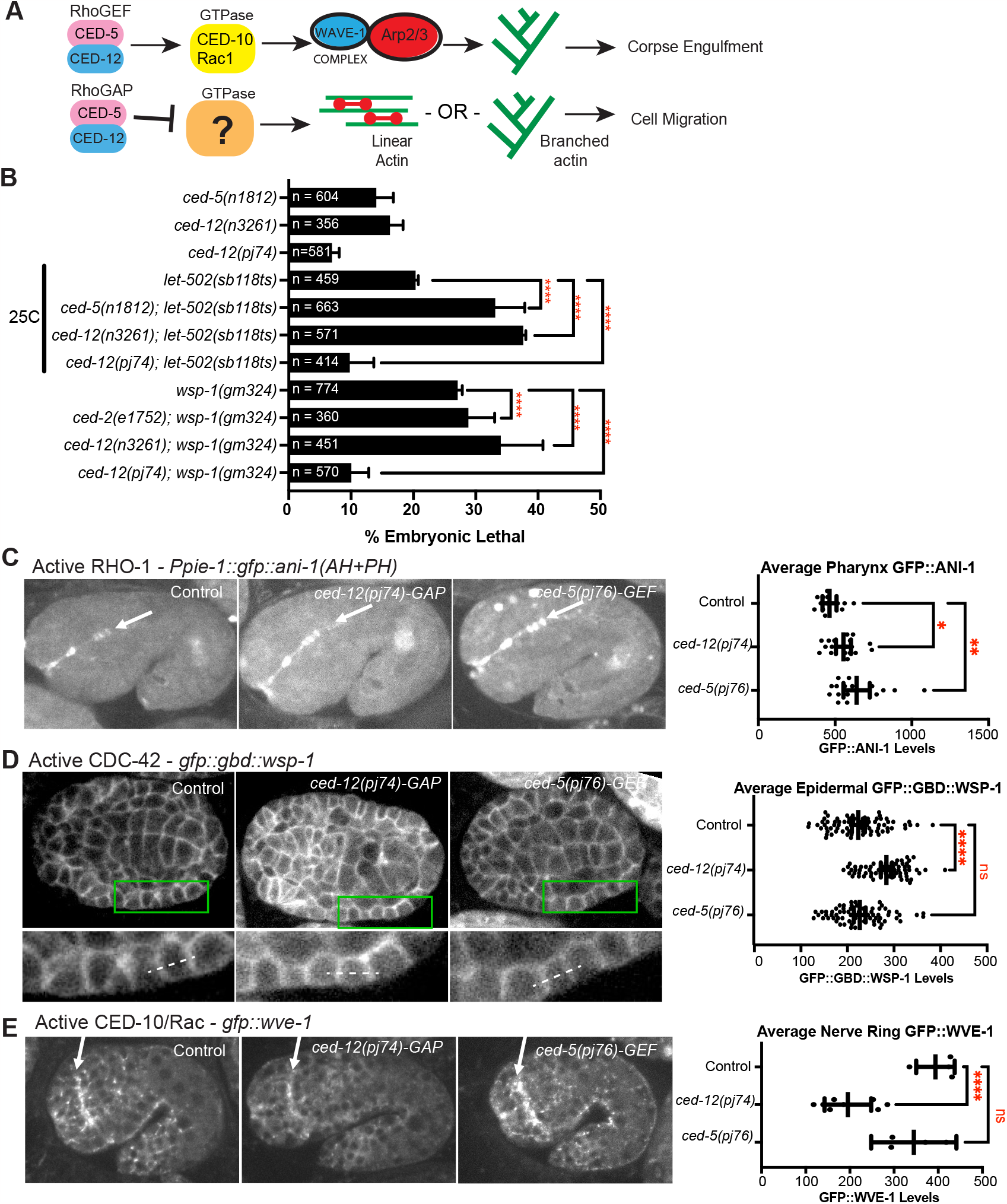
Evidence that the GAP function of CED-5/CED-12 affects RHO-1/RhoA and CDC-42. (A) While the CED-5/CED-12 GEF function is known to active the GTPase CED-10/Rac1 during corpse engulfment, the GTPase target of the proposed GAP function of CED-5/CED-12, proposed to regulate cell migration, is not known. (B) Genetic analysis compared embryonic lethality of *ced-5* and *ced-12* single mutants and double mutants with the RHO-1 pathway, using Rho Kinase mutant, *let-502(sb118ts)*, and with the CDC-42 pathway, using WASP mutant *wsp-1(gm324). let-502* experiments were done at 25C since this is a temperature sensitive allele (Vanneste et al. 2013). See also Table 3. (C) A biosensor for active RHO-1, *Ppie-1::gfp::ani-1(AH+PH)* (Tse et al., 2012), was used to compare the effects of mutating the proposed GAP and GEF regions on CED-12 and CED-5. Levels of active RHO-1 at the apical pharynx of 1.5 fold embryos (white arow) were measured. Embryos are oriented with anterior to the left and dorsal up. (D) A biosensor for active CDC-42, *gfp::gbd::wsp-1* (Kumfer et al., 2010), imaged from a ventral view, anterior at left., allowed comparison of active CDC-42 at cell junctions in lateral epidermal cells, at approximately 320 minutes after first cleavage, 23 C. (E) Since a Rac biosensor is not available we used the Rac GTPase target, gfp::wve-1 (REF) to compare “Active Rac” levels. The signal at the nerve ring (white arrow) was compared. Embryos are oriented anterior to the left and dorsal up, at the 1.5 fold stage. Asterisks denote statistical significance, *=p<0.05, **=p<0.01, ****=p<0.0001, as determined by one-way Welch’s ANOVA test.

To specifically mutate the GEF domain of CED-5/CED-12, we used the structure alignment based on DOCK2/ELMO1 (Chang et al., 2020) and consulted the DOCK5/ELMO1 structure (Kukimoto-Niino et al., 2021). DOCK2 has a helix predicted to make multiple interactions with Rac1, required for promoting its activation, and sometimes referred to as the Rac binding domain (RBD). CED-5 has a similar helix, with conserved amino acids, like DOCK2 S1528, conserved in CED-5 as S1541 (Fig. 4B-C). M1529 of DOCK2 is replaced in CED-5 by R1542. Using UCSF Chimera (Pettersen et al, 2004; Goddard et al., 2018), we predicted that R1542 would make similar hydrogen bonds with Rac W56/CED-10 W56. Therefore, to specifically interfere with the GEF activity, we used CRISPR to mutate S1541 and R1542 to Alanines. This mutant, *ced-5(p76)* SR1541/1542AA, had similar embryonic lethality as the *ced-5* or *ced-12* null alleles (13%, Table 1).

**Table 3.** Effects of CED-5/CED-12 GAP mutation on RHO-1 and CDC-42 pathways.

| Genotype | n | %Lethality | Standard deviation |
| --- | --- | --- | --- |
| *rho-1 RNAi* | 293 | 0.14 | 0.0045 |
| *ced-5(n1812); rho-1 RNAi* | 313 | 0.14 | 0.03 |
| *ced-12(n3261); rho-1 RNAi* | 324 | 0.28 | 0.017 |
| *ced-12(pj74); rho-1 RNAi* | 418 | 0.76 | 0.02 |
| *cdc-42 RNAi* | 537 | 0.51 | 0.17 |
| *ced-5(n1812); cdc-42 RNAi* | 350 | 0.38 | 0.21 |
| *ced-12(n3261); cdc-42 RNAi* | 227 | 0.43 | 0.085 |
| *ced-12(pj74); cdc-42 RNAi* | 417 | 0.11 | 0.079 |
| *let-502(sb118ts) - 25C* | 459 | 0.20 | 0.005 |
| *ced-5(n1812); let-502(sb118ts) - 25C* | 663 | 0.33 | 0.048 |
| *ced-12(n3261); let-502(sb118ts) - 25C* | 571 | 0.38 | 0.0049 |
| *ced-12(pj74); let-502(sb118ts) - 25C* | 414 | 0.10 | 0.039 |
| *wsp-1(gm324)* | 774 | 0.27 | 0.008 |
| *ced-2(e1752); wsp-1(gm324)* | 360 | 0.29 | 0.04 |
| *ced-12(n3261); wsp-1(gm324)* | 451 | 0.34 | 0.07 |
| *ced-12(pj74); wsp-1(gm324)* | 570 | 0.10 | 0.029 |

### Evidence that CED-5/CED-12 regulate two other GTPases, RHO-1/RhoA and CDC-42

The CED-12 GAP motif, when threaded with the structure of DOCK2/ELMO1 bound to active Rac1 (Chang et al., 2020), is oriented away from the Rac binding interface (Fig. 4B). This suggested that the CED-12/CED-5 GAP activity may have other GTPase targets other than Rac/CED-10 (Fig. 5A). A genetic test used in *C. elegans* to identify GTPase targets is to cross the candidate GAP to hypomorphic alleles, or partial loss of function, of the target GTPases. The prediction is that loss of a GAP in combination with a hypomorphic allele will rescue the loss of function phenotype (Neukomm et al., 2011). In contrast, we predicted that loss of a GEF in combination with a hypomorphic allele will synergistically enhance the loss of function phenotype. Therefore, we combined partial loss of function mutations for the three main *C. elegans* GTPases, *ced-10/Rac, cdc-42* and *rho-1*, with either loss of *ced-5 (n1812*), loss of *ced-12 (n3261*), or the CRISPR mutations generated based on the structure, *ced-12* GAP *(pj74)*. We predicted that loss of GAP function would suppress partial loss of a GTPase.

To test for interactions with RHO-1/Rho, we used a temperature sensitive mutation in Rho Kinase, *let-502(sb1008ts) (*Vanneste et al., 2013). At 25 C *let-502(sb1008ts)* resulted in 20% embryonic lethality, and in combination with null alleles of *ced-5 or ced-12*, this rose to 33% and 37%, respectively, while in combination with the *ced-12* GAP allele this dropped to 10% (Fig. 5B). Similarly, RNAi of *rho-1*, fed to control worms for two days, resulted in 14% embryonic lethality, and fed to *ced-5 or ced-12* null worms, led to 14% and 28% embryonic lethality, respectively, while depleting *rho-1* in *ced-12(pj74)* GAP mutant reduced lethality to 8%, and this drop was significant (Table 3). These results suggest the GAP activity of CED-12 acts on RHO-1, while the GEF function acts in parallel to RHO-1.

To test for interactions with CDC-42, we used mutations in the CDC-42 effector WSP-1. The null allele *wsp-1(gm324)* showed 27% embryonic lethality, and combining *wsp-1* with *ced-12(n32561 null*) led to 41% (n=451), while combining *wsp-1* with the *ced-12* GAP mutant reduced lethality to 10% (n=570) (Fig. 5B). Similarly, *cdc-42 RNAi* after 2 days led to 51% embryonic lethality, that was not significantly changed by complete loss of *ced-5* or *ced-12*, but was suppressed significantly by loss of the GAP function (10% lethal) (Table 3). Thus, the GAP activity of CED-12 may be acting on CDC-42.

#### Effects on levels of active RHO-1/RhoA

Active RHO-1 in embryos can be measured with a biosensor that uses a region of ANI-1/anillin that binds to active RHO-1, *pie-1p::gfp::ani-1(AH+PH*) (Tse et al., 2012). Loss of the GAP activity, using *ced-12(pj74)* resulted in a significant increase in levels of active RHO-1, as seen by the increase signal at the apical pharynx, and buccal cavity (Fig. 5C). Loss of the proposed GEF function, using *ced-5(p76)* resulted in even more elevated active RHO-1. This result supported the genetic result that the GAP function regulates RHO-1.

#### Effects on levels of active CDC-42

Active CDC-42 in embryos can be measured with a biosensor that uses the GTPase binding domain of WSP-1, *cdc-42p::gfp::GBD::wsp-1* (Kumfer et al., 2010; Zilberman et al., 2017). Loss of GAP activity using *ced-12(pj74)* resulted in a significant increase in levels of epidermal *cdc-42p::gfp::GBD::wsp-1*, whereas loss of GEF activity through *ced-5(pj76)* resulted in no significant change (Fig. 5D). This result supported the genetic result that the GAP function regulates CDC-42.

#### Effects on levels of active Rac1/CED-10

Since a Rac biosensor does not exist for monitoring embryos, we instead measured effects on the Rac effector, GFP::WVE-1. Active Rac1/CED-10 is required to activate the WAVE complex (Miki et al, 1998). Neither loss of the GAP function, nor the GEF function increased the levels of GFP::WVE-1. Instead, loss of the GAP function using *ced-12(pj74)* resulted in significantly reduced *gfp::wve-1* levels in all tissues measured, including in the nerve ring (Fig. 5E). *ced-5(p76)* resulted in no significant change. This result further supported that the GEF function does not simply promote active Rac, and suggested CED-12 is not a GAP for CED-10/Rac1.

Altogether these genetic and gene expression studies strongly support that the GAP activity of CED-5/CED-12 can regulate the GTPases RHO-1 and CDC-42.

### CED-12 GAP (R537A) allele affects epidermal migrations, not corpse engulfment

To test if the embryonic lethality in the *ced-12(pj74)* R537A proposed GAP mutant correlated with changes in F-actin at the leading edge of migrating epidermal cells, we crossed this mutant into the epidermal F-actin strain (*lin-26p::LAmCherry*), and found that F-actin levels increased, similarly to null alleles in *ced-12* (Fig. 6A,B). Examining the corpses demonstrated that *ced-12(pj74)* R537A had normal corpse engulfment, and wild-type levels of F-actin around corpses, in contrast to the abnormal corpse engulfment and low levels of F-actin around corpses seen in the *ced-12(n3261)* null allele (Fig. 6A,C-D). These results suggested that CED-12 has a GAP function that is required for epidermal cell migrations of ventral enclosure, but dispensable for corpse engulfment (Fig. 6D,E).

**Fig. 6:**
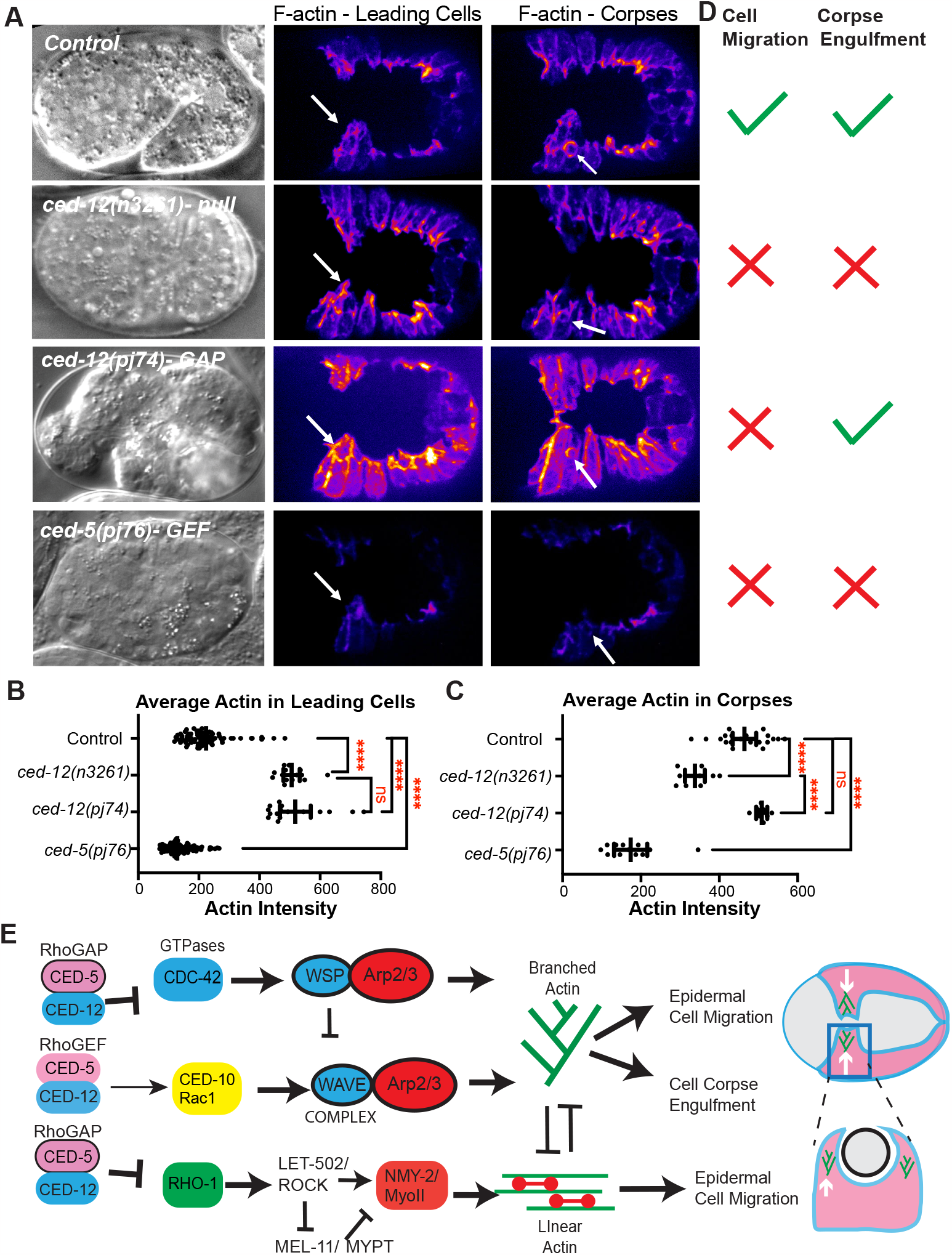
CED-12 GAP mutant alters cell migration, not corpse engulfment, while CED-5 Rac binding domain mutant alters both. (A) Embryos were imaged by DIC (left column) and using *lin-26p::LifeAct::mCherry* to visualize cell migration defects, corpse engulfment defects and F-actin levels. F-actin is shown during ventral enclosure. Center column white arrows pointing to LCs. Right column white arrows indicate C2 corpses in the epidermis. (B) Average intensity of F-actin in LCs during ventral enclosure was analyzed as in Fig. 1. (C) Average intensity of F-actin in C1, C2, and C3 corpses was measured as in Fig. 3. (D) Summary of the phenotypes, regarding defects in epidermal cell migration, or in corpse engulfment, for each genotype. Asterisks denote statistical significance, ****=p<0.0001, as determined by one-way Welch’s ANOVA test. (E) Model summarizing CED-5/CED-12’s role as a GEF towards CED-10 during corpse and engulfment, and a GAP towards CDC-42 and RHO-1 during cell migration. The embryonic lethality and active GTPase levels contributed by mutations in the CED-5/CED-12 proposed GAP function (R357A) and proposed GEF function (SR1541/1542AA) (Fig. 5) and the effects on epidermal and corpse F-actin (Fig. 6A-D) leads us to propose this model: The GEF function, as expected, regulates CED-10/WAVE to support corpse engulfment. However, during epidermal cell migration, the role of CED-5/CED-12 is better explained as the result of the GAP function regulating branched and linear actin through the GTPases CDC-42 and RHO-1. Together these results illustrate how the bifunctional CED-5/CED-12 complex can support embryonic development through a bifunctional role on GTPase regulation.

### CED-5 GEF (SR1541-1542AA) affects corpses, and epidermal migrations

The *ced-5(p76)* mutation, a two amino acid change in the Rac binding domain, predicted to block GEF activity (Fig. 4B-C), affected corpse engulfment, with persistent unengulfed corpses, and highly reduced F-actin around corpses (Fig. 6A,C), as would be expected if this mutation alters the GEF function of CED-5/CED-12. In the migrating epidermal cells, *ced-5(p76)* significantly reduced F-actin, a distinct phenotype compared to complete loss of CED-12 *(n3261)* or the CED-12 GAP mutant *(pj74)*, which both increase epidermal F-actin (Fig. 6A,B,D). Therefore, while mutating the CED-12 GAP function only affected the cell migration function of CED-5/CED-12, and not the corpse engulfment function, mutating the GEF function in CED-5 affected both processes.

## DISCUSSION

### CED-5/CED-12 have previously undescribed role in embryonic morphogenesis

The discovery of CED-12 was impactful, since it identified a molecule, conserved across phyla, that affected cell migrations and cell engulfment of apoptotic corpses (Gumienny et al. 2001; Wu et al, 2001; Chung et al, 2000; Zhou et al., 2001). The cell migration defects referred to the larval migrations of the distal tip cell (DTC). These early papers hinted at a possible embryonic role (Gumienny et al., 2001). However, the proposed embryonic role had not been further explored. Our findings here definitively show that the embryonic lethality of *ced-12*, also seen in partner protein *ced-5*, is due to failure to regulate F-actin during embryonic cell migrations, resulting in increased F-actin, altered dynamics and faster epidermal enclosure as the cells increase their migration rate.

The reason this embryonic role was mostly missed is that even with null alleles, the penetrance is relatively low, with only 12% of embryos dying (Table 1). This is in contrast to 100% embryonic lethality seen with complete loss of Rac1/CED-10, or loss of WAVE or Arp2/3 complex components. Loss of *ced-5* or *ced-12* leads to lower lethality, and high instead of low F-actin in the migrating epidermis. These differences suggest two conclusions: (1) there is likely a second pathway, in addition to CED-5/CED-12, that regulates epidermal F-actin during this migration and (2) too much F-actin in the epidermal cells disrupts migrations. The *ced-5* and *ced-12* embryonic phenotype, with fully penetrant increased F-actin and partially penetrant embryonic lethality (Fig. 1, 6, Table 1) has been shown for other mutations in proteins that regulate actin. For example, loss of *hum-7/Myo9*, leads to increased F-actin in all embryos, and 6% embryonic lethality (Wallace et al., 2018).

Our results here showed additional roles for CED-5/CED-12 during embryonic development. Mutant *ced-5* and *ced-12* embryos were often shorter than normal, and reduced length correlated with reduced viability (Fig. 2). Wild-type embryos are cuboidal, but once they enter the spermatheca and are fertilized, they are squeezed into an oval shape. The force for this may come from actomyosin in the spermatheca. Mutations in the *rho-1* pathway, like filamin, or the Rho GAP SPV-1 lead to embryos that are round (Kovacevic and Cram, 2010; Tan and Zaidel-Bar, 2015.), so too much or too little actomyosin contractility can alter egg shape, including producing round embryos. Our results suggest that at these earliest stages of embryonic development, when oocytes are being fertilized, the branched actin regulators CED-5/CED-12, CED-10 and WAVE complex components, are required so that embryos are the correct length and shape, by promoting the correct levels of RHO-1 activity.

Null mutations in CED-5 or CED-12 have consistently higher levels of F-actin in the migrating epidermal cells, and this correlated with faster migrations, and increased dynamics in the protrusions and retractions at the edge of leading cells (Fig. 1). Surprisingly, the same epidermal cells that showed increased F-actin in the migrating cells, also show reduced F-actin around corpses (Fig. 3). To untangle this puzzling dual nature of the CED-5/CED-12 GEF, we searched for new functions, and discovered them by comparing CED-12 to other ELMOs and ELMODs, and also to other GAP proteins. It has been shown that ELMODs mainly function as GAPs for Arf GTPases in processes such as protein traffic (Ivanova et al., 2014). We identified a clear GAP domain within CED-12, that better aligned to ELMODs, not ELMOs, which do not contain a GAP domain (Fig. 4). Therefore, we tested if *C. elegans* uses CED-12 as a bifunctional ELMO/ELMOD.

We used the recently published structures of DOCK2/ELMO1/Rac1 and DOCK5/ELMO1/Rac1 to help us test the bifunctional hypothesis for CED-12 function. One idea that we considered, based on the structure, was that CED-12 may have two roles, depending on the open vs. closed conformation for the entire complex. For example, while the GAP residues point away from bound active Rac1, perhaps in the closed conformation CED-12 could help hydrolyze active Rac1 to its GDP state. Based on the published structures, this is not likely. The catalytic Arginine, R537, points away from the Rac binding domain in both open and closed conformations. This outward orientation of R537 actually increases the interactions that CED-12 can make, since the GAP motif is available in both open and closed conformations to interact with additional proteins. This led us to test if CED-12 could couple the activation of one GTPase with the turn-over of a second GTPase.

We provide evidence that two other GTPases are likely targets of the CED-12 GAP function. Genetic analysis and measurements of levels of biosensors for active GTPases suggested that CED-12 GAP function regulates RHO-1 and CDC-42 (Fig. 5). Since the effect includes rescuing embryonic lethality, this result suggests that excess levels of CDC-42 and RHO-1 are detrimental for embryonic epidermal morphogenesis. Our studies of other proteins that when missing result in dead embryos due to excess epidermal F-actin levels are in agreement with this finding that excess F-actin must be controlled for healthy cell migrations (Bernadskaya et al., 2012, Wallace et al. 2018, Raduwan et al., 2020).

We tried to devise surgical changes that would only affect the GEF function, similar to how we removed the GAP function. This was more complicated. Our first attempt to mutate the GEF function was to mutate the PXXPXXP domain at the C terminus of CED-12, proposed to support GEF function (Gumienny et al., 2001; Lu et al., 2005). This mutation led to low level embryonic lethality (4%) and no change in epidermal F-actin levels (Table 1, data not shown). We next tried to interfere with GEF function by mutating two residues that mediate binding to Rac1, based on the structures. This mutant, *ced-5(pj76)*, resembled null mutations in *ced-5* and *ced-12*, since it altered corpse engulfment and epidermal cell migration, with similar levels of embryonic lethality. However, it had opposite effects on epidermal F-actin levels (lowering them) (Fig. 6A,B). One explanation for why mutating these two Rac1 binding residues is worse than simply deleting *ced-5* is that this “GEF” mutation traps active Rac1 molecules, acting as a Rac1 sink. We have proposed that other GEFs can activate Rac1, but in animals with the *ced-5(pj76)* mutations, other Rac1 GEFs may lose access to Rac1. Interestingly, this mutant also led to increased active RHO-1 (Fig. 5C).

To incorporate all of these findings, we propose the following model for CED-5/CED-12 function (Fig. 5E). To properly promote branched actin to engulf corpses, the GEF function of CED-5/CED-12 activates CED-10/Rac1, exactly as has previously been proposed. To regulate F-actin during embryonic epidermal cell migration, a different GEF must work to activate CED-10. There are 18 other GEFs found in *C. elegans*. Several have been shown to regulate embryonic development, mainly by regulating CDC-42 or RHO-1 (Motegi and Sugimoto, 2006, Kumfer et al., 2010, Chan et al., 2015). We are investigating GEFs that act on CED-10.

During epidermal cell migrations, CED-5/CED-12 have an important role limiting the activity of CDC-42, and this requires the proposed CED-12 GAP function. We previously published the surprising observation that in *wsp-1(gm324)* mutants, which are approximately 30% lethal, epidermal F-actin is too high, and the cells migrate too quickly (Raduwan et al., 2020). This resembles what we show here for loss of the CED-12 GAP function. Perhaps in GAP mutants with excess CDC-42, too much WSP-1 is activated, which also promotes excessive branched F-actin, which is detrimental. This suggests too much or too little WSP-1 has same phenotype: too much F-actin, and some dead embryos.

During epidermal cell migrations, CED-5/CED-12 have an important role limiting the activity of RHO-1, and this requires the proposed CED-12 GAP function (Fig. 5B,C, Fig. 6E), and may also require the CED-5/CED-12 GEF function (Fig. 5C, Fig. 6A,B, E). If RHO-1 is overly active, more actomyosin would form, driving more linear F-actin. It was possible that excess RHO-1 would increase membrane and cortical tension, thus inhibiting protrusions, as has been proposed (Diz-Munoz et al., 2013). However, increased epidermal F-actin correlated with increased leading edge membrane dynamics (Fig. 1), suggesting excess RHO-1 supported protrusions, perhaps through increased tension, as has been in some migrating cells (Reviewed in Sens and Plastino 2015). A second consequence of excess RHO-1, given that levels of actin can be limiting in some cell types (Suarez and Kovar 2016), is reduced monomeric actin available to form branched actin. At this time, we do not have tools to specifically monitor branched actin vs. linear actin in these cells, so this prediction that excess RHO-1 leads to excess linear actin at the expense of branched actin, will have to await new tools.

Attempts to tag CED-12 with GFP using CRISPR are underway. When we can image endogenous CED-12 in live animals, a strong prediction is that there will be different populations of CED-12, carrying out distinct functions, both promoting and inhibiting F-actin, in different parts of the epidermal cells.

## Materials and Methods

### C. elegans strains built for this paper

OX764 *ced-5(n1812); lin-26p::LifeAct::mCherry* ; OX869 *ced-12(n3261); lin-26p::LifeAct::mCherry* ; OX924 *ced-10(tm597)/nT1-gfp; lin-26p::LifeAct::mCherry* ; OX799 *ced-5(n1812); lin-26p::LifeAct::mCherry; dlg-1::gfp* ; OX778 *ced-12(k149); dlg-1::gfp* ; OX775 *ced-(e1752); dlg-1::gfp* ; OX293 *ced-(tm597)/nT1-gfp; dlg-1::gfp* ; OX994 *lin-26p::LifeAct::mCherry; smIs34[ced-1p::ced-1::gfp::rol-6(su1006)]* ; OX1002 *ced-5(n1812); lin-26p::LifeAct::mCherry; smIs34[ced-1p::ced-1::gfp::rol-6(su1006)]* ; OX1003 *ced-12(n3261); lin-26p::LifeAct::mCherry; smIs34[ced-1p::ced-1::gfp::rol-6(su1006)]* ; OX989 *ced-12(pj74)* ; OX1026 *ced-5(pj76)* ; OX991 *ced-12(pj74); lin-26p::LifeAct::mCherry; dlg-1::gfp* ; OX1046 *ced-5(pj76); lin-26p::LifeAct::mCherry; smIs34[ced-1p::ced-1::gfp::rol-6(su1006)]* ; OX1047 *ced-5(n1812); let-502(sb118ts)* ; OX1048 *ced-12(n3261); let-502(sb118ts)* ; OX1049 *ced-12(pj74); let-502(sb118ts)* ; OX1050 *ced-2(e1752); wsp-1(gm324)* ; OX1051 *ced-12(n3261); wsp-1(gm324)* ; OX1052 *ced-12(pj74); wsp-1(gm324)* ; OX1040 *ced-5(pj76); gfp::wve-1* ; OX1041 *ced-12(pj74); gfp::wve-1* ; OX1042 *ced-5(pj76); cdc-42p::gfp::gbd::wsp-1* ; OX1043 *ced-12(pj74); cdc-42p::gfp::gbd::wsp-1* ; OX1044 *ced-5(pj76); pie-1p::gfp::ani-1(AH+PH)* ; OX1045 *ced-12(pj74); pie-1p::gfp::ani-1(AH+PH)* ; OX776 *ced-2(e1752); ced-12(k149)*

### Strains used in this paper

JUP38 *lin-26p::LifeAct::mCherry* ; FT48 *xnIs16[dlg-1::gfp]* ; OX669 *pj64[gfp::3xflag::wve-1]* ; FT1459 *xnIs506 [cdc-42p::gbd-wsp-1::gfp]* ; MG617 *xsSi5[pie-1p::gfp::ani-1(AH+PH)::pie-1 3’UTR + Cbr-unc-119(+)]* ; HR1157 *let-502(sb118ts)* ; NG324 *wsp-1(gm324)* ; MT4434 *ced-5(n1812)* ; MT11068 *ced-12(n3261)* ; NF87 *ced-12(k149)* ; CB3257 *ced-2(e1752)* ; MT5013 *ced-10(n1993)* ; LE198 *ced-10(tm597)/nT1-gfp*.

### RNAi experiments

All RNAi bacterial strains used in this study were administered by the feeding protocol as in (Sasidharan et al., 2018). RNAi feeding experiments were done at 23°C unless otherwise mentioned. Worms were synchronized and transferred onto seeded plate containing RNAi-expressing bacteria. To monitor effectiveness of the RNAi we used two methods. We counted the percent dead embryos, which after two days is expected at >90% for *gex-3*. We also monitored post-embryonic silencing of a gfp-tagged strain in the intestine, such as *gfp::gex-3*. All RNAi treatments were done for three days.

### Live Imaging

Imaging was done in a temperature-controlled room set to 23°C on a Laser Spinning Disk Confocal Microscope with a Yokogawa CSUX scan head, on a Zeiss AxioImager Z1 Microscope using the Plan-Apo 63X/1.4NA or Plan-Apo 40X/1.3NA oil lenses. Images were captured on a Hamamatsu CMOS Camera using MetaMorph software, and analyzed using ImageJ. Controls and mutants were imaged within 3 days of each other with the same imaging conditions. All measurements were performed on raw data using ImageJ. For fluorescent measurements, background intensity was subtracted by using a box or line of the same size and measuring average intensity in the same focal plane, near the animal.

### Quantitation of immunofluorescence

Quantitation of live fluorescence was performed using the line selection and the dynamic profile function of ImageJ to measure fluorescence along lines of equal lengths. For all experiments shown, the images were captured at the same exposure settings for wild type and mutants. All quantitation was done on the raw images. The Fig. legends indicate when images were enhanced for contrast, and the same enhancement was applied to a mosaic of the related images for that experiment. Each measurement was taken following the subtraction of background fluorescence.

### Statistical Analysis

For grouped data in Fig. 1, statistical significance was established by performing a one-way Analysis of Variance (ANOVA), the Brown-Forysythe and Welch ANOVA, followed by a Dunnett’s multiple comparisons T3 post-test. For ungrouped data in other Fig.s, an unpaired t-test, the unequal variance (Welch) t test, was used. Error bars show 95% confidence intervals. Asterisks (*) denote p values *= p<.05, ** = p<0.001, *** = p<0.0001, ***=p<0.00001. All statistical analysis was performed using GraphPad Prism 8.

*UCSF Chimera*: https://www.rbvi.ucsf.edu/chimerax_ModelingCED-5/CED-12 with and without CED-10: CED-5/CED-12 were modeled bound to CED-10 and in its apo form using DOCK2/ELMO1/Rac1 and DOCK2/ELMO1 coordinates0020(PDB IDs: 6TGC and 6TGB). The models were generated using SWISS-MODEL, and visualized in UCSF Chimera.

## Acknowledgements

We thank the NCRR-funded *Caenorhabditis* Genetics center (CGC), funded by NIH Office of Research Infrastructure Programs (P40 OD010440), for strains. Molecular graphics and analyses performed with UCSF Chimera, developed by the Resource for Biocomputing, Visualization, and Informatics at the University of California, San Francisco, with support from NIH P41-GM103311. We thank members of the Soto Lab and William Wadsworth for comments. This research was funded by a grant from the National Institutes of Health (NIH) (GM081670) to M.C.S., and used a Spinning Disk Microscope acquired through an NIH Shared Instrumentation Grant (1S10OD010572) to M.C.S.

## Notes

### Competing Interest Statement

The authors have declared no competing interest.

